# Catamers: Multi-specific therapeutics that concatenate individual warheads on a DNA scaffold via Watson-Crick interactions

**DOI:** 10.64898/2026.09.23.753027

**Authors:** Jasper Kamb, Shimin Xu, Roland Bürli, Jiajia Cui, Han Xu, Alexander Kamb

**Affiliations:** Catessa Biotherapeutics, 321 Dedalera Dr., Portola Valley, CA 94028

**Keywords:** Nucleic acid therapeutic, DNA scaffold, ADC, pinocytosis, modular design

## Abstract

We describe the concept for a new modality to create multi-specific therapeutics that utilizes modified nucleic acids as a scaffold to concatenate multiple warheads (hence called “catamers”). Catamers are assembled via Watson-Crick interactions into multi-specific molecules from individual components such as ligand-binding modules, protease-cleavage sites, and toxins. Each warhead is covalently bound to an oligonucleotide, which in turn is connected to the catamer scaffold. Post-synthesis of the individual catamer components, they are annealed together such that Watson-Crick base pairs on the oligonucleotides dictate the organization of the warheads on the complete catamer. Notably, catamers are *not* aptamers, though they can contain aptamers as warheads, along with small molecules and peptides. By design, catamers are intended to address three key problems that have hamstrung the field of multi-specifics: (i) optimization rate; (ii) manufacturing cost; and (iii) unpredictable immunogenicity (Amash et al., 2024 PMID: 39189686). Catamers are purely synthetic and do not require cultured cells to produce. These advantages are based on the underlying design of the catamer and will require experimental validation in future studies. As a proof-of-concept catamer, we focus on a prostate-specific membrane antigen (PSMA)-targeted toxin conjugate. In molecular simulations, this catamer displays structural characteristics to support its intended pharmacology. Importantly, the design addresses one of the key shortcomings of antibody drug-conjugates (ADCs): off-target toxicity caused by release of the toxin after pinocytosis by healthy cells in contact with the blood.

**Graphical abstract:** Model of a catamer drug-conjugate with functional moieties comprised of a double stranded DNA scaffold whose complementary strands are connected to a PSMA-binding peptidic moiety (green) and, on the other strand, to three different functional groups linked together: (i) a toxin (deruxtecan, red); (ii) a PSMA protease-cleavage site (yellow); and (iii) a half-life extending moiety (myristate, magenta).

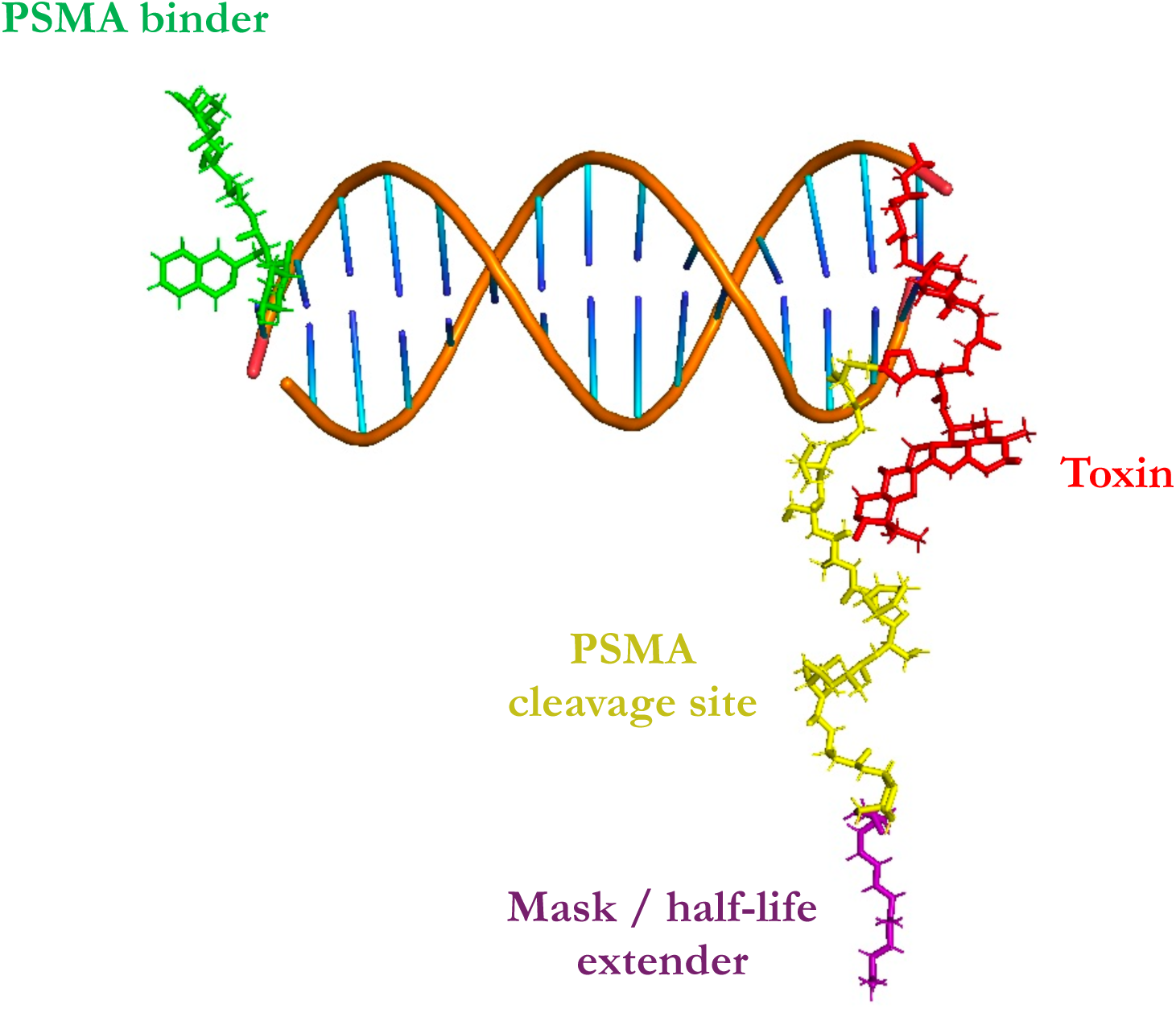

## INTRODUCTION

The importance of multi-specific molecules in modern drug discovery cannot be overstated. In an era of increasingly limited options for medicines that bind a single target, a compelling case can be made to combine pharmacological activities into multi-specifics that can integrate and coordinate binding to multiple antigens. Novel combinations of binding and functional elements have produced bispecific agents that solve certain therapeutic problems with an elegant simplicity. One example is Hemlibra^TM^, a bispecific antibody that mimics activated factor 8a (FVIIIa) in hemophilia A patients who lack the F8 gene, by bridging coagulation factor IXa and X (FIXa and X), and orienting them to activate the downstream clotting process^1^. Another impressive example of bispecifics is the T cell engager Blincyto^TM^ which redirects the cytotoxicity of T cells to CD19^+^ blood cancers^2^. A third example is antibody-drug conjugates, a kind of multi-specific therapeutic devised in the 1990s that may have reached its apotheosis with the breast cancer drug Enhertu^TM3,4^.

Despite their utility, recombinant multi-specific proteins face several challenges that are likely to become more problematic as poly-functionality is increased. Proteins evolved to replace other molecular types as the chief effectors of biological processes, most notably small molecules and nucleic acids. These other biological molecules have significant limitations with regard to modularity and diversity, respectively. Low-molecular-weight molecules are subject to effects that are difficult to predict and non-additive at this small scale. Electronegativity, resonance and other quantum mechanical features become dominant, hindering accurate prediction of chemical behavior of merged structures from the properties of their individual substituents. Despite these challenges, proteins will continue to occupy an important place in the arsenal of therapeutic modalities. Nucleic acids on the other hand, though much improved in diversity through synthetic modifications, are fundamentally constrained by certain structural features such as phosphate charge and base stacking. Further, their distribution within the body following systemic administration is often limited and it can be challenging to deliver nucleic acids to the intracellular compartment of a target cell in a specific organ or a tumor.

Proteins also have considerable limitations as multi-specific therapeutic agents. Though the modularity and chemical diversity of polypeptides was arguably the key innovation in the evolution of life once replication was achieved, non-natural, complex multi-domain proteins are often difficult to make in the laboratory, let alone manufacture at scale. Enormous proteins exist (e.g., titin), but these evolved over hundreds of millions of years. It is not easy to splice together multiple domains into a single functional unit that folds efficiently and retains the desirable pharmacological function. Drug discoverers struggle to find general protein formats that can be utilized for complex designs and scaled for manufacturing. Iterative optimization cycle times can be hampered by the difficulty of expressing and purifying sufficient amounts of homogeneous material for testing. Finally, a product that meets pharmacological and manufacturing specifications can still fail in the clinic due to adaptive immune response, a property that has proven impossible to predict reliably. Could some of these problems be addressed using a different template for multi-functionality?

Nucleic acids are a type of polymer that have exceptional advantages in this context; namely, very predictable structures that are determined largely by Watson-Crick base pairing. The predictable geometry of the double helix, and its easily estimated stability as a function of length and sequence, are huge advantages, especially in the era of computation and machine learning. We therefore selected DNA as the basis for a new scaffold to replace proteins. The idea is to use a modified DNA single strand as a scaffold to organize complementary oligonucleotides (oligos) connected to warheads which anneal in an orderly way to the scaffold to form the final concatenated, noncovalent complex, or catamer (Fig. 1A). Use of modified bases and backbones addresses previous issues associated with nucleic acid therapeutics; for example, susceptibility to nuclease degradation and innate immune reactivity. Past decades of work by chemists focused on RNAi, antisense, and aptamers have created an inventory of modifications that overcome these formidable obstacles^5,6^. In addition, by using warheads that are not displayed via major histocompatibility antigens to T cells (i.e., small molecules, peptides < 9 residues, and aptamers), catamers evade a potential adaptive immune response.

**Fig. 1:**
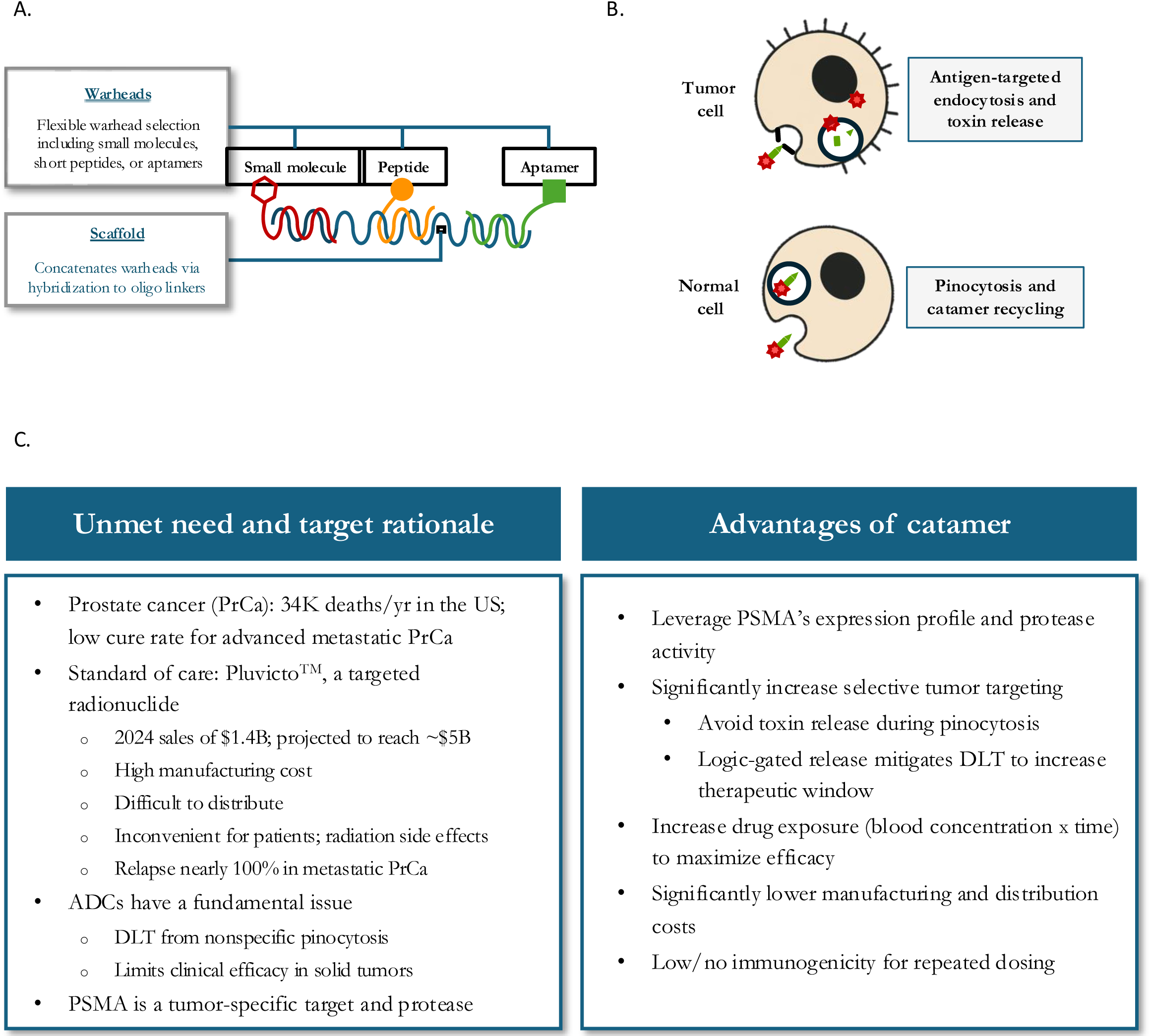
General composition of catamers and a use case (toxin conjugates). **A.** Molecular composition of catamers. Catamers contain multiple warheads connected via a nucleic acid scaffold. Examples of multi-specifics include drug-conjugates, T-cell engagers, and multi-specific modulators (antagonists and agonists, or both). Shown is a diagram of a catamer with a scaffold that binds 3 complementary oligos, each linked to a different type of warhead. **B.** Sequence of steps for a drug-conjugate catamer mechanism. In the case described in the text, the catamer consists of a toxin conjugated to a PSMA-binding moiety or warhead (red star). The catamer is designed to traffic to the lysosome and release toxin only when it binds to a PSMA-expressing tumor cell (see text for details). If the catamer is internalized nonspecifically via pinocytosis, it is recycled to the surface and released from the cell in an intact form. **C.** Summary of rationale for a PSMA catamer compared to other modalities.

Here we undertake simulation of one specific catamer design, based on warhead components that have been tested in the clinic. The catamer is directed at PSMA, a validated target for prostate cancer, and is designed to deliver a toxin to tumor cells that express the PSMA antigen. Thus, it is the catamer analog of an antibody-drug conjugate. This construct also incorporates features intended to address one of the most problematic aspects of ADCs: unscheduled, off-target release of toxin in the blood compartment. We argue that the catamer platform holds promise as a new multi-specific modality with a high degree of modularity. The modular, ‘Lego-like’ composition allows catamers to be rapidly optimized for a specific therapeutic purpose with great structural and functional precision.

## RESULTS

### Overall design of catamer and intended pharmacology

The molecular architecture of the catamer takes advantage of two proteases, PSMA and prostate specific antigen (PSA), that are well known to be expressed in healthy prostate tissues and prostate cancers but show limited expression in other tissues. Because of the strong validation of PSMA as a target, we began with the goal of creating a catamer drug-conjugate that incorporates a PSMA-binding warhead and a half-life-extending molecule (e.g., myristate) that binds human serum albumin (HSA). As part of its normal function in cells, neonatal Fc receptor (FcRN) binds HSA (as well as the Fc portion of antibodies) to allow HSA to recycle when engulfed by the cell in endosomes, thus extending its half-life^7^. By co-opting the HSA recycling mechanism, catamer half-life is expected to increase. In addition, this mechanism prevents toxin release unless the catamer is bound to a cancer cell via its PSMA-binding warhead. When PSMA is bound to the catamer, the protease-cleavage site is hydrolyzed by a membrane-associated protease (PSMA or PSA in this case), thereby releasing myristate. Catamer cleavage by one of these proteases not only prevents recycling to the surface but also triggers the release of the toxin via cellular proteases and/or nucleases. Toxin release only occurs when a membrane-proximate protease such as PSMA or PSA cleaves the half-life-extending moiety, setting in train a series of events that lead to creation of a free, membrane-permeant toxin.

Various warheads were included in the catamer to accomplish these mechanistic objectives (Fig. 1B, C). The overall design was based on the following intended pharmacology in a PSMA-expressing tumor cell:

1. The catamer binds to PSMA target on the tumor cell
2. The catamer is retained on the surface or internalized via endocytosis
3. PSA (or PSMA, depending on which recognition sequence is used in the catamer) hydrolyses the peptide imbedded in the linker sequence, removing the half-life extender to reveal a “soft spot” to RNAses and/or proteases inside or outside the cell
4. The toxin is released by protease/RNAse cleavage
5. The toxin diffuses into the cytoplasm of the cell and initiates apoptosis

In contrast to tumor-cell pharmacology, the catamer was designed for the following pharmacology in non-malignant, PSMA-negative cells:

1. The catamer is internalized via non-specific pinocytosis
2. In the absence of binding to PSMA, the PSA recognition sequence imbedded in the catamer is uncleaved and the catamer remains attached to HSA via its myristate moiety
3. The intact catamer recycles back to the extracellular milieu

### Warheads with covalently attached oligos

The first warhead was designed to bind the catamer to PSMA on the tumor cell surface via a peptide mimetic of Pluvicto^TM^ ^8–10^, a clinically effective prostate cancer medicine (Fig. 2A). The peptide warhead was linked to a 15-mer oligo of arbitrary sequence with 8 G/Cs and 7 A/Ts (see Methods). Of the many possibilities for attachment, we chose an amide bond between the warhead amine and the 5’ position of the terminal nucleotide (Fig. 2B).

**Fig. 2:**
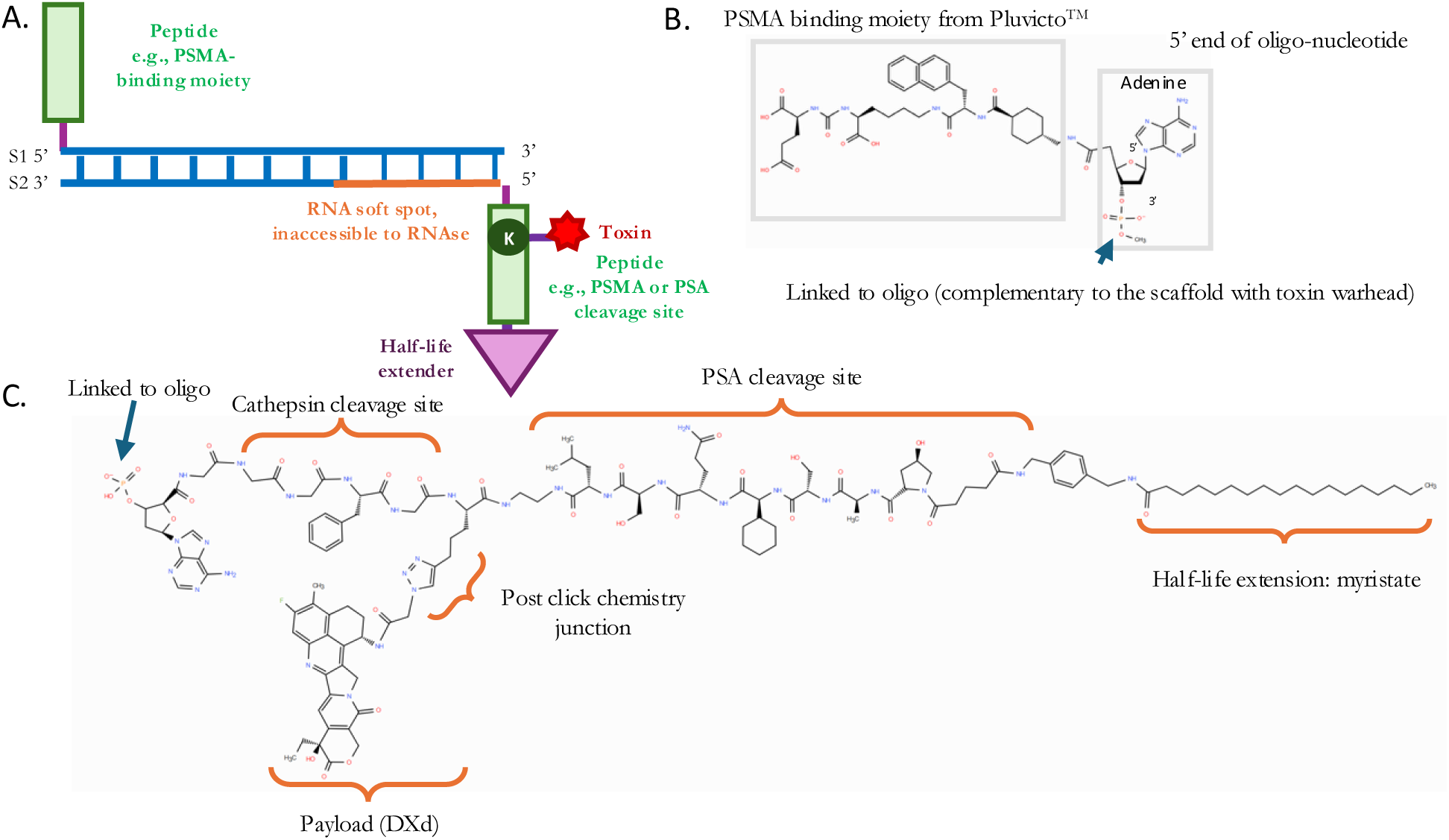
Catamer design and warheads. **A.** Diagram of PSMA drug-conjugate design. Protease-cleavage sites (e.g., for PSMA or PSA) are incorporated to release the half-life-extender (e.g., myristate) and then free toxin (e.g., deruxtecan (DXd), SN38, etc.) in the lysosome where it is trimmed of its associated amino acids by cathepsins to form a molecule that kills the cell when it diffuses out of the lysosome, in a manner similar to the mechanism of toxin release of the ADC Enhertu^TM^. The DNA portion of the catamer is depicted with a blue ladder. This depiction is illustrative; for actual bp numbers, see text. **B.** PSMA-binding warhead and linker used in the modeling. 5’ and 3’ positions on the ribose ring are labeled. **C.** Toxin warhead (DXd) with PSA cleavage site and other functionality used in the modeling.

We next designed the more complicated, multi-functional warhead located at the other end of the catamer, opposite the PSMA-binding warhead. Myristate was attached to a PSA- or PSMA-specific protease-cleavage site via an amide bond^11–13^. The purpose of this module is to permit the FcRN-recycling mechanism to operate absent the PSA (or PSMA; see Fig. S2) protease, enforcing a requirement for two events to release the toxin: PSMA ligand-binding and PSA protease cleavage to free the myristate moiety from the catamer.

More proximal to the DNA scaffold, a toxin (deruxtecan (Dxd) or SN38) was linked to a variant of a cathepsin cleavage site (Fig. 2C). This sequence is known to result in the release and trimming of Dxd to a membrane-permeant form^14^. Dxd and SN38 were attached via a triazole linkage, the product of certain click-chemistry reactions^15^. Note that with catamers, other covalent couplings and release mechanisms are possible; for example, RNA/DNA “soft spots” vulnerable to cellular nucleases rather than cathepsin cleavage sites, or a combination thereof, to release the active toxin. Finally, the multi-component warhead was attached to a second 15-mer oligo complementary in sequence to the first one. When annealed, these two warhead-linked oligos form the final PSMA toxin-conjugate catamer.

### Simulation of catamer molecular interactions

The PSMA catamer was modeled in three dimensions using molecular dynamics (MD) simulations. The basic catamer was built from its components and energy-minimized first without the protein-binding partners (Fig. 3A). As expected, the 15-mer DNA helix was above 350 K, well above body temperature and the temperature used for modeling (300 K). Thus, it is predicted to remain stable in the bloodstream even though the principal catamer warheads, the PSMA binder and the toxin, are not covalently to each other; rather, they are connected via Watson-Crick base pairs. The length and composition of this oligo pair can be readily varied to further increase stability if needed.

**Fig. 3:**
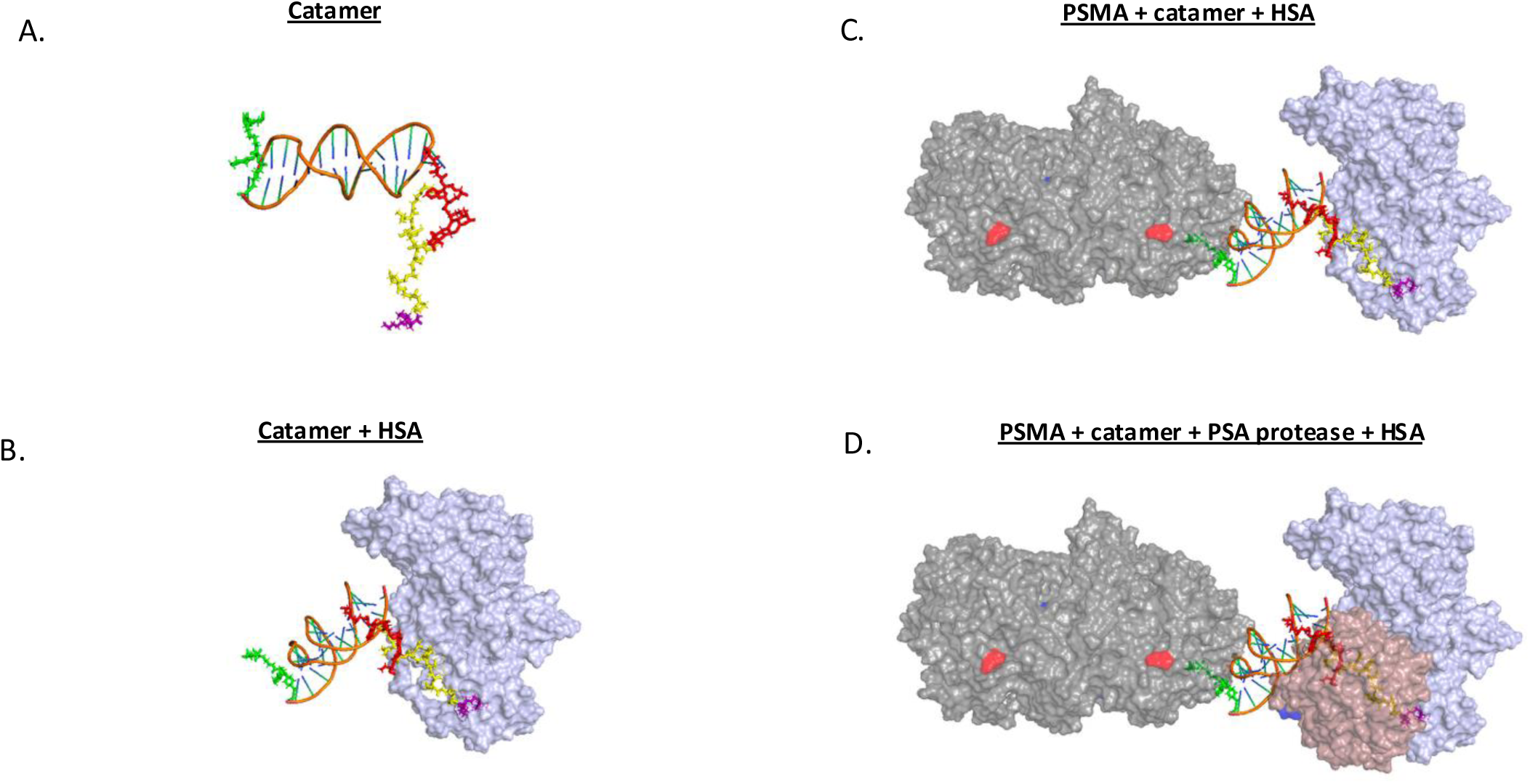
Catamer structures modeled. **A.** Catamer structure without binding partners, oriented in extended form to clarify the scaffold and warheads. Myristate is magenta; DXd is yellow; the PSA cleavage site is red; and the PSMA-binding warhead is green. Energy-minimized catamer structures bound: HSA (purple). **B.** Energy minimized structures bound: HSA+PSMA (grey). **C.** Energy-minimized structures bound: HSA+PSMA+PSA protease (brown). The PSMA molecule C-terminus is labeled in red (small dot on the grey structure). The N-terminus is not visible but is pointing down, presumably toward the cell membrane (PSMA is a type II membrane protein).

The protein-binding partners of each warhead were added sequentially and energy-minimized to produce intermediate structures. First, the catamer with HSA bound was modeled (Fig. 3B). This structure simulated the situation where the catamer is bound to a tumor cell prior to dissolution of the toxin conjugate. To ensure that FcRN did not interfere with binding of HSA to the catamer, we modeled the HSA/FcRN complex from its cocrystal structure^16^ (Fig. S3B). The FcRN bound on the other side of the HSA protein, away from the catamer. Next the PSMA extracellular domain was added to the complex (Fig. 3C). Because PSMA is a type II membrane protein, the globular portion was oriented with its N-terminus, pointed down towards the membrane. Finally, a model of the PSA protease was included with the other components of the catamer complex to simulate attack on the cleavage site proximal to myristate, an event that breaks the cycle of ingestion via pinocytosis and return to the membrane surface (Fig. 3D). Following this proteolysis, the catamer remnant traffics to the lysosome, where proteases and/or nucleases liberate the toxin, which decomposes into a membrane-permeant topoisomerase I inhibitor. Minimization via MD revealed that when the peptide linker was situated correctly in the PSA active site, the two proteins (HSA and PSA) were quite close, with their van der Waals surfaces within 5 Å. This problem was exacerbated when a PSMA cleavage site was substituted within the catamer for PSA cleavage site, a situation designed to exploit a second PSMA molecule on the cell surface to provide the activity that cleaves the myristate from the catamer, triggering the series of steps leading to toxin release in the tumor cell. In the cases where the PSMA protease was used, the catamer contained SN38 toxin to illustrate the generality of the catamer design. However, because the PSMA protease is significantly larger, substitution of the PSMA protease for PSA resulted in a considerable steric clash (Fig. S3). This suggested the value of testing catamers where the spacing between the myristate and PSA cleavage sequence was increased.

### Optimization of PSMA catamer

To optimize the distance between the myristate and protease recognition sequences, polyether linkers (i.e., spacers) were added and after visual inspection of different lengths, a [CHO]_6_ polyether was selected for energy minimization (Fig. 4A). Each link added an average of ∼1.8 Å to the spacing. The final linker yielded a molecular complex with better access of the PSA protease to its cleavage site. The efficiency of the cleavage process measured empirically will ultimately determine what linker spacing should be used, but simulations suggest a length in the range of a few Å may be optimal for the PSA protease complex. Notably, changing the linker to the other polymeric units, e.g., 5 or 20 glycines, seemed to have little effect on the energetics of the system, suggesting tolerance to a variety of linker chemistries [Fig. S4B,C; Table S4). The complex with the larger PSMA protease substituted for PSA required a longer spacer, a [CHO]_20_ was used in the simulation (Fig. 4B). Although the scaffold was optimized *in silico*, additional optimization of the nucleic acid chemistry may be required to maintain scaffold integrity *in vivo*.

**Fig. 4:**
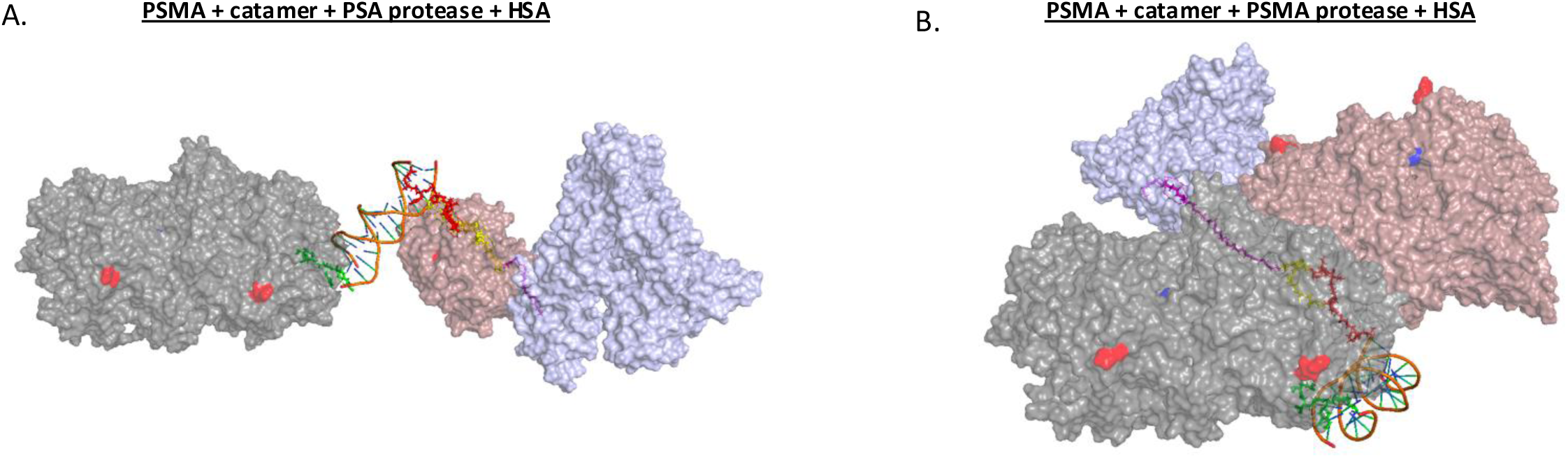
Models with catamers that contain a longer linker between the myristate and protease cleavage sites. The protease and HSA are separated by 5-6 Å in both cases. **A.** Energy-minimized model with 6 [COH] units (polyether) between the PSA protease cleavage site and myristate, with the PSMA protease bound (brown). **B.** Similar complex with the PSMA protease bound in place of PSA and energy-minimized, but the linker/spacer is 20 [COH] units. The complex is shown at a slightly different scale. A different view, more clearly revealing the individual proteins is shown in Fig. S4A. The red dot on the grey structure is the PSMA C-terminus. The membrane is predicted to be below the grey PSMA molecule (C-terminus of PSMA visible, colored with red).

### Addition of a second antigen-binding domain to the catamer

Antigen escape is a common problem for effective immuno-therapeutics, including ADCs^17^. This is especially true if the therapy does not target an essential function (such as EGFR). For PSMA-targeted therapies, resistance to the radionuclide Pluvicto^TM^ has been associated with antigen loss^18^.

One readily available mitigation is to combine two targeting agents as has been common practice in oncology and infectious disease, notably with HIV drugs^19^. In the case of catamers, it is straightforward to add a second antigen-targeting moiety by extending the catamer scaffold slightly to accommodate another warhead that binds an antigen other than PSMA on prostate cancer cells. Integrins are seldom perfectly tissue-specific, but integrin αvβ6 (ITGV/ITGB6) is recognized as relatively prostate-specific, compared to other integrins^20^. Because the proteases that release the toxin of the catamer are prostate-specific, it is reasonable to use this integrin as a second targeting antigen. In addition, both high affinity peptides and small molecules that bind αvβ6 exist^21,22^.

We selected one of these, the peptide NAVPNLRGDLQVLAQKVART (A20FMDV2) derived from foot-and-mouth disease virus. This peptide is a known selective inhibitor of αvβ6 with an IC50 of 3 nM, ∼1000 fold more specific over other RGD-directed integrins^23^ (e.g., αvβ3, αvβ5, and α5β1). Though this peptide is perhaps twice the length desired for catamers to minimize the chances of immune response, it served the purpose to illustrate the ease of increasing the complexity of catamer designs. Because a cocrystal structure between integrin and the A20FMDV2 peptide was not available, we used AlphaFold to generate a model and checked it using a cocrystal structure between a minibinder and αvβ6 thought to bind at the same site^24^ (Fig. S5).

To design the dual-antigen-binding catamer, one of the oligos (the “scaffold”) was extended to 30 residues and used to guide assembly of two scaffold-complementary 15-mer oligos with attached warheads (Fig. 5). Two formats were considered. One (Format A) contains the antigens at the opposite ends of the catamer (Fig. 5A). The other (Format B) has one antigen (PSMA) at the end and the other (integrin) near the middle (Fig. 5B). The PSMA-binding warhead was attached as before. For the integrin-binding warhead (A20FMDV2), the peptide was attached at the 5’ end of either its guide oligo (Format A) or the 3’ end of its guide oligo (Format B) via its N-terminus using an amide bond (see Methods). This N-terminal portion of the peptide has been shown to be non-essential for binding to the integrin^25,26^. Again, the oligo sequence was arbitrary, with 8 G/Cs and 7 A/Ts, the same composition as the other oligos. The oligos were virtually annealed to the 30-mer scaffold such that the two 15-mers, separated by a nick, formed a double helix with the complementary 30-mer scaffold.

**Fig. 5:**
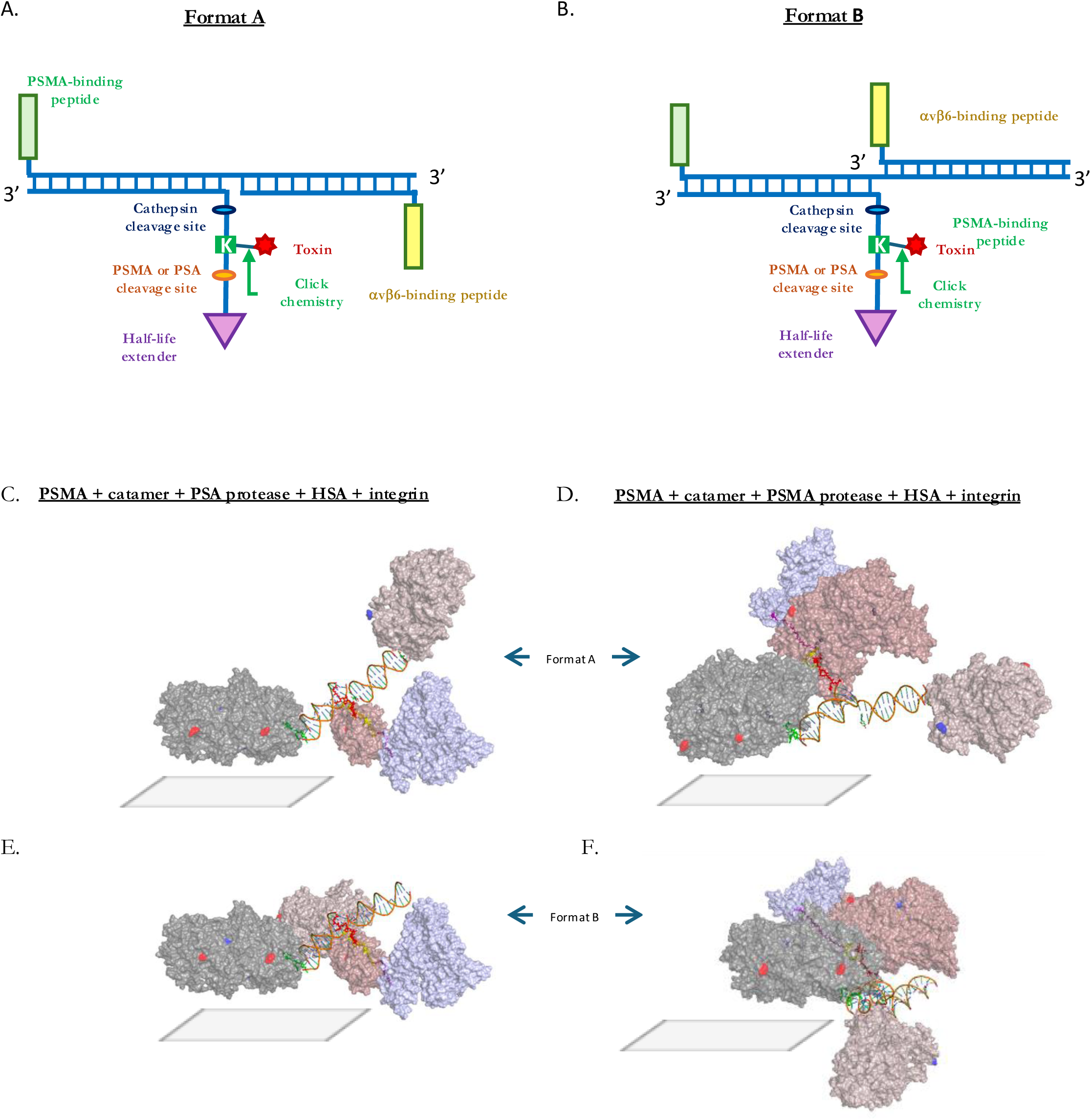
Diagram of two versions of a PSMA drug-conjugate with an extension to add a second warhead peptide that binds integrin avb6. In the case modeled here, the scaffold catamer is a 30-mer and the two complementary oligos are 15-mers. Protease-cleavage sites (e.g., for PSMA or PSA) are incorporated to release the half-life-extender (e.g., myristate) and then free toxin in the lysosome where it is trimmed of its associated amino acids. The blue ladder, depicting the DNA double helix, is illustrative; for actual bp numbers, see text. **A.** Format A: two antigen-binding warheads placed at ends of the scaffold. **B.** Format B: two antigen-binding warheads placed closer on the scaffold by linking to the 3’ end of one oligo (avb6) warhead) and the 5’ end of the other (PSMA warhead**). C-F.** Energy-minimized molecular complex of catamer with 30-mer scaffold and 2 warheads attached via complementary 15-mer oligos, one of which binds PSMA and the other of which binds avb6 (light brown). The C-termini are labeled in red and the N-termini, where visible, in blue. Format A structures (as in Fig. 5A) are in the top two panels (A and D); Format B structures are in the bottom panels (E and F). The grey parallelograms represent the predicted plane of the cell membrane based on the orientation of the termini of PSMA and integrin**. C.** PSA (brown) bound to cleavage site (brown) with 6 units of polyether spacer. Left panel has standard orientation of PSMA surface antigen with membrane below the complex. **D.** PSMA (brown) bound to cleavage site with 20 units of polyether spacer. Structure is rotated to give a better view of the integrin. Note the bend in DNA at the nick in the double strand. **E.** Same proteins bound as in (C) by a Format B catamer. **F.** Same proteins bound as in (D) by a Format B catamer.

This entire catamer was rebuilt with the different binding proteins attached, including human αvβ6, and subjected to MD energy minimization. The polyether linkers were included in this simulation as above. The resultant structure suggested simultaneous occupancy of all the catamer binding partners, including the PSA protease, was possible (Fig. 5C). The structures, with the polyether linker of 20 units, accommodated the large PSMA protease as well (Fig. 5D).

Not surprisingly, the situation was different when Format B was modeled. The complex with the PSA protease was reasonable ((Fig. 5E,S5). However, the PSMA protease was fairly crowded near the nick in the DNA component (Fig. 5F). Perhaps more importantly, the orientation of the PSMA and integrin antigens (as well as HSA) with respect to the predicted location of the cell membrane, was potentially problematic. We therefore returned to the question of optimizing the catamer complex with special attention to membrane orientation.

### Optimization of PSMA/integrin catamer

To modify the catamers and reposition its geometry relative to the membrane, we took advantage of the highly predictable structure of the DNA helix. We altered the catamer backbone in two ways. First, we added an additional 5 bp of double-stranded structure by extending the PSMA warhead oligo 5 bases and adding 5 bp complementary to the scaffold (Fig. 6A). This resulted in rotation of the relative positions of HSA and PSMA by roughly 180° due to the pitch of the double helix, mitigating concerns that HSA-binding might interfere with PSMA-binding at the membrane surface (Fig 6B,C). Second, we explored the effect of adding 5 bases of extra single-stranded structure to the scaffold. To eliminate any problems caused by the hydrophobic nature of the bases, only the phosphodiester backbone was included, with a hydrogen atom in place of the base at the 1’ position. This approach provides further flexibility to the overall catamer structure in case of unfavorable steric interactions at the membrane surface. Both these modifications produced reasonable structures when energy-minimized. The model with 5 extra bp of double-stranded region was especially clear with respect to optimal positioning of the protein components. The other complex with 5 unpaired bases was slightly skewed regarding the relative positions of the proteins and membrane surface, but the extra degrees of freedom in the middle of the catamer backbone likely ameliorate this potential issue, allowing avidity to come into play on the membrane (Fig. 6D).

**Fig. 6:**
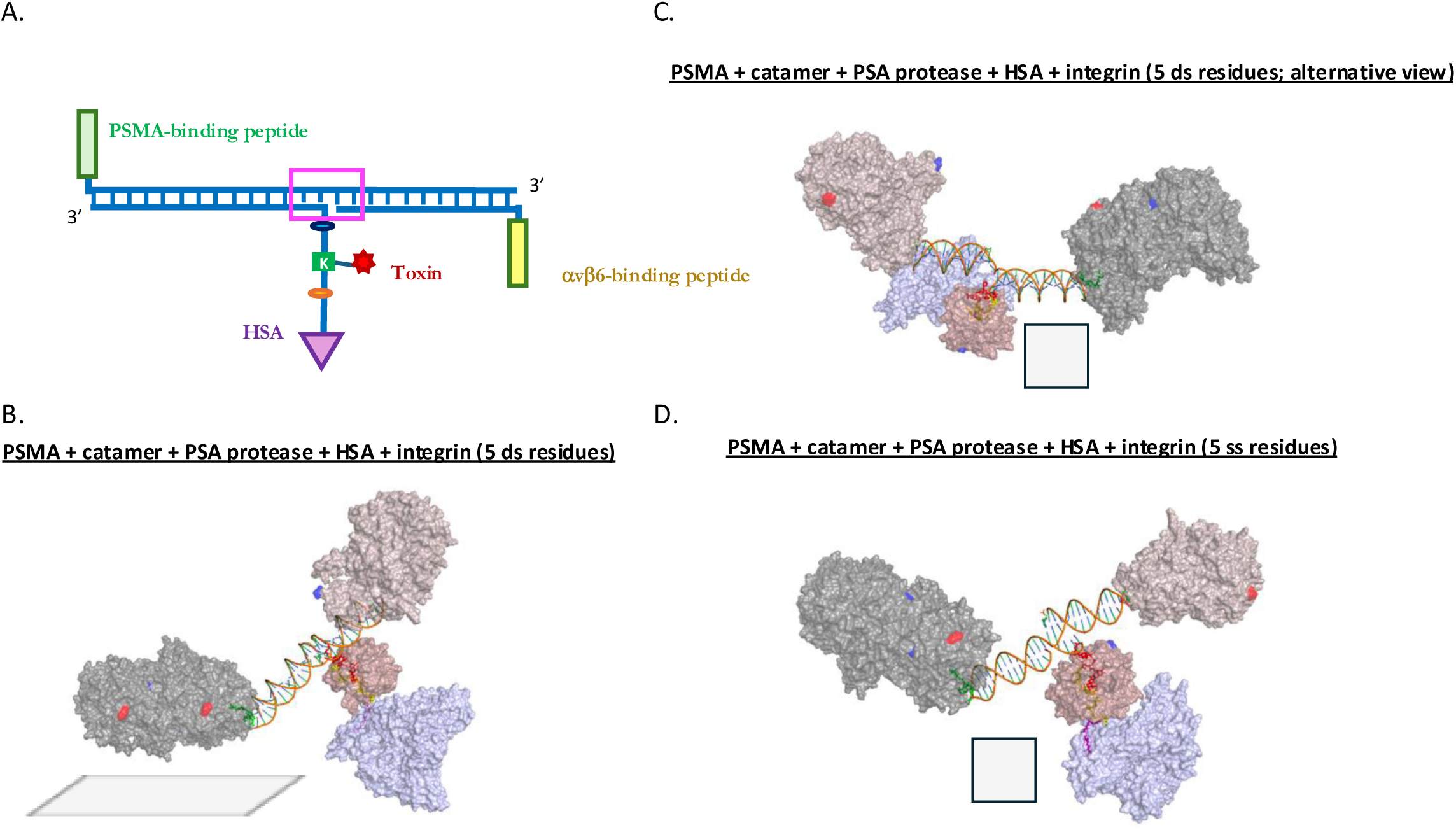
Catamer structures with 5 residues added to extend the DNA scaffold. **A.** Diagram showing the position of the extra 5 residues, boxed in magenta. The 5 bases are modeled as either double-stranded (ds) or a gap with 5 DNA backbone linkages with a hydrogen in place of the base to form a pseudo-single-stranded regions (ss), depending on the catamer structure (see text). In this diagram, only the ds version is shown. **B.** Energy-minimized complex with linker (spacer) of 6 [CHO] units. The orientation is chosen to emphasize the location of the membrane (down and perpendicular to the plane of the image, indicated by the grey parallelogram). **C.** Alternative view showing the N- and C-termini of the antigens. In this orientation the membrane is predicted to be below the complex in the same plane as the image (perpendicular to reader), illustrated by the grey square. **D.** Catamer complex with 5 residues added to the scaffold to form a ss region between the ds catamer ends (see Methods). Grey square represents the membrane plane, perpendicular to reader.

## DISCUSSION

Because of the finite coding capacity of the human genome, known for nearly 25 years, few new single targets are likely to be uncovered in the future. The drug discovery field, which has depended so much on delineation of new targets, now approaches an asymptote where most single protein targets discovered via germline genetics, somatic cell genetics, etc. have been identified^27^. Powerful as it is, AI cannot overcome the fact that the human genome has only ∼20,000 genes. Thus, in our view it is imperative that new ways of using known targets are explored. An obvious approach is multi-specifics that can target or bridge together different disease-relevant proteins and functional modules^28^.

We considered the possibility that polymers other than polypeptides might be able to solve the problem of configuring functional modules or warheads with the same precision but greater tractability in a synthetic setting. Most proteins, even those encoded by bacteria, are much larger than might be expected based on the ligands they interact with. For example, β-galactosidase is a tetramer of subunits that are over 1,000 amino acids, despite catalyzing the simple hydrolysis of lactose. Antibodies are roughly 150 kDa, though their binding regions comprise a small fraction of this total mass. Much of the antibody protein structure serves as a scaffold to orient binding sites optimally. True modularity of design was not apparently necessary to evolve complex life forms with their associated protein-based molecular machinery. But in the context of drug discovery, the challenges are different. There is no universal modular system for multi-specific proteins, and to date, many bispecifics do not work in a standard format and most be tested empirically in multiple formats, of which there are dozens^29^. Partly due to the challenges of biosynthesis, multi-specific “frankenmolecules” are difficult and time consuming to optimize and problematic to manufacture. Finally, protein multi-specifics elicit adaptive immunity at a discouraging rate, blunting efficacy and causing trial termination, especially for repeat subcutaneous dosing^30^.

Catamers take advantage of predictable, simple chemistry of nucleic acids, including Watson-Crick interactions. They are the therapeutic version of DNA origami, a field that has demonstrated beautifully the power of nucleic acids to form Lego-like assemblies^31^. A notable example is the use of Holliday junction structures to attach warheads^32^. Other ways of creating branched structures for the scaffold may also be envisioned^33^. Catamers are purely synthetic; cell culture is not involved in their production. Catamers (scaffold, oligos, warheads, and binding proteins) can be readily simulated and their geometry optimized by varying scaffold sequence length and composition. In addition, the catamer backbone and bases can be modified as illustrated in the work described here. Stability of nucleic acids, a major stumbling block in the initial phase of nucleic acid therapeutics, has been overcome by incorporation of appropriate modifications to DNA and RNA. Nearly 20 nucleic acid therapeutics, mostly DNA antisense and siRNA, are clinically approved^34^. In one extreme example of stability, a DNA molecule composed of L-nucleotides was stable for a year in pond water while the same standard B-DNA sequence disappeared in hours^35^.

The catamer example described here incorporates two prostate-selective proteases, one of which (PSA) is secreted. However, in the body PSA is rapidly bound by serpins and other serine protease inhibitors that essentially eliminate enzymatic activity^36^. Thus, the tumor selectivity of the PSMA catamer depends on localization at the surface of prostate cancer cells that express high levels of PSA, and cleavage by the secreted PSA before inhibition occurs. This mechanism has been exploited previously by a prodrug (L-377,202) developed by Merck over two decades ago^12^. This investigational medicine had the expected pharmacology in human clinical trials but was discontinued^13^.

As an alternative, we also explored the use of PSMA, a membrane-anchored metalloprotease, as the distal release mechanism of the toxin. Though in some cancer patients PSMA can be shed from the surface into the bloodstream, soluble PSMA is thought to be largely inactive. Whether or not the level of active PSMA is sufficient to cleave the alternative catamer is not clear. The stability of the two catamer types (PSA vs. PSMA cleavage-dependent) could be readily tested in patients’ blood and cell-based assays to select a candidate with superior properties.

The focus here was on PSMA as a cancer target, and the warhead modules and covalent linkers were selected among many possible as the basis for modeling a toxin-conjugate. However, much broader applications of the catamer modality are possible. Other ADCs, T-cell engagers, and multi-specific designs are possible, from simple catamers that include a tissue-localization module to more complex designs that incorporate so-called logic-gates by analogy with binary computational devices^37^ (e.g., OR, AND, NOT). Warhead choices are only limited by the compatibility with the chemistry of nucleic acids and the desire to maintain low immunogenicity. Thus, small molecules, peptides, and aptamers are available. The technologies for isolation of such warhead binders have advanced tremendously over the last 2-3 decades: (i) phage display for peptides^38^; (ii) DNA-encoded libraries for small molecules and peptides^39,40^ (which come pre-designed with an oligo attached); and (iii) SELEX for aptamers^41^. Note also that catamers lend themselves to warheads that are not ultra-high affinity because multiple copies can be readily incorporated to exploit avidity.

Catamers do not solve all problems associated with drug discovery. Most significantly, they are confined by the same set of targets encoded by the human genome as protein modalities. Like proteins, they are not suited to diffusion across membranes in the way that certain small molecules can.

## CONCLUSIONS

Catamers are a new modality that may complement traditional protein-based multi-specifics, including T-cell engagers and antibody-drug conjugates. Their modular construction via Watson-Crick base-pairing offers several unique advantages. A precise and predictable assembly of multiple functional elements to a well-defined structural template allows optimization of the catamer in a rapid manner and offers great versatility in exchanging and optimizing the functional warheads. As such, new molecular warheads can be added with minimal effort, like another bead to a string. Importantly, catamers are produced by chemical synthesis rather than assembled from (or within) cultured cells. Compared to preparation of many multi-specific engineered proteins manufactured to date, this process is likely to result in lower costs, better yields and higher structural accuracy. Finally, catamers are uniquely suited for chronic treatments that must avoid an adaptive immune response.

## METHODS

### Molecule building

To create the DNA scaffold, the core sequence 5’-ATACCAGCTTATTCA-3’ was used with the 3D-DART software package^42^. B-form DNA was specified; otherwise, default parameters from the 3D-DART server (accessed via Docker) were used. To make the longer catamer scaffold, this core sequence was duplicated for the main scaffold strand, while the reverse complement of the core sequences was used for both warheads.

Warhead molecules, constructed with SMILES strings, were read via the rdkit library (RDKit: Open-source cheminformatics. https://www.rdkit.org) so that Python scripts could be used for the visualization of molecules in the Jupyter Notebook environment, before saving as .pdb files. The following SMILES strings were used for the different catamer warhead components:

1. NH2 DA5 = "Nc1ncnc2n(cnc12)[C@@H]3O[C@H](CN)[C@@H](OP(=O)([O-])[O-])[C@H]3"
2. COOH DT5 = "CC1=CN(C(=O)NC1=O)C2CC(C(O2)C(=O)O)OP(=O)(O)O"
3. Half-life extender (myristate) = "CCCCCCCCCCCCCCC(=O)[O-]"
4. PEG-linked half-life extender = "CCCCCCCCCCCCCCCCCOC(=O)N"
5. Glycine myristate = "CCCCCCCCCCCCCC(=O)NCC(=O)N"
6. PSMA binder (Pluvicto binder moiety) = "C1CC(CCC1CNC(=O)CO)C(=O)N[C@@H](Cc1cc2ccccc2cc1)C(=O)NCCCC(C(=O)O) NC(=O)N[C@@H](CCC(=O)O)C(=O)O"
7. Toxin for PSA (deruxtecan) = "NCC(NCC(NCC(N[C@@H](CC2=CC=CC=C2)C(NCC(NCOC(N=[N+]=[N-])C(N[C@@H]3C4=C5C(C(N6C5)=CC([C@](O)(C(OC7)=O)CC)=C7C6=O)NC8=CC(F)=C(C)C(CC3)=C48)=O)=O)=O)=O)=O)=O"
8. Toxin for PSMA (SN-38) = "CC(NCC(N[C@@H](CC1=CC=CC=C1)C(NCC(N[C@@H](CCCC2=CN(CCCO[C@]3(CC)C(OCC4=C3C=C5N(CC6=C(CC)C7=CC(O)=CC=C7N=C65)C4=O)=O)N=N2)C(N CC)=O)=O)=O)=O)=O”
9. PSA cleavage site = "C#CCCC[C@@H]C(=O)NCCNC(=O)[C@@H](CC(C)C)NC(=O)[C@H](CO)NC(=O)[C@@H](C[C@H]C(=O)N)NC(=O)[C@@H](C1CCCCC1)NC(=O)[C@@H](CO)NC(=O)[C@H](C)NC(=O)C1C[C@@H](O)CN1C(=O)CCCC(=O)NCC1=CC=CC=C1CN"
10. PSMA cleavage site = "NC(=O)CC(N)C(=O)NC([C@@](=O)O)CCC(=O)NC([C@](=O)O)CCC(=O)NC([C@ @](=O)O)CCC(=O)NC([C@](=O)O)CCC(=O)NCC"

As each molecule was built, PyMOL was used to determine atom numbering for bonding^43^. Peptides, other warhead moieties, and the DNA scaffold were loaded using the command line version of LEaP (Assisted Model Building with Energy Refinement) to build, parameterize, and solvate the final catamer. To connect the warheads and oligos, terminal atoms at linkage sites were removed, enabling covalent bond formation. Atom types were adjusted where required to maintain GAFF2 compatibility, such that covalent bonds could be formed to connect peptides to the scaffold. ParmEd and cpptraj were used to inspect .parm7 and .rst7 files to verify atom names, numbers, types, bonds, charges, etc.

For the spacing optimization phase, units of either glycine or ethers were added to myristate via Python scripts as described above prior to minimization.

### Energy minimization of catamer

All minimization was performed using Amber^44^ with explicit water. To parameterize peptides for minimization, Antechamber was used to calculate partial charges with a GAFF2 forcefield and AM1-BCC (Austin Model 1 with Bond Charge Corrections^45^). Missing bonded or nonbonded parameters like bond, angle, dihedral, and vdW were assigned with parmchk2. Initially, the catamer was restrained to allow approximate solvation, followed by a final step of unrestrained minimization of the whole system.

Heat was added from 0 to 300 K under constant volume (NVT). Density equilibration was executed under constant pressure (NPT) at 1 atm. Final production molecular dynamics was performed under constant temperature (300 K) and pressure (1 atm). Energy terms were checked and the structure was inspected to ensure acceptable features.

### Addition of proteins and minimization of catamer complexes

Once the catamer was built and energy-minimized, the catamer-binding proteins (HSA, PSMA, PSA, αvβ6 integrin) were added to .pdb files using the same steps described above. The HSA .pdb file was downloaded from https://www.rcsb.org/structure/1AO6^46^; the PSMA .pdb file from https://www.rcsb.org/structure/1Z8L^47^ and the PSMA active sites were determined. The PSA .pdb structural coordinates were downloaded from https://www.rcsb.org/structure/2ZCK^48^. The integrin structural coordinates were obtained from https://www.rcsb.org/structure/8TCG^24^ and the minibinder portion of the co-crystal complex was removed from the file and replaced with the A20FMDV2 peptide generated via AlphaFold3^49^. FcRN was downloaded from https://www.rcsb.org/structure/4N0F, the bound beta-2-microglobulin and HSA were removed and then the isolated FcRN was manually placed for visualization to interact with the catamer’s HSA via PyMOL.

Energy minimization of complexes was performed by the same set of steps described above for the catamer alone.

The additional 5 bases were (TAGTA) added to the 3’ strand (strand with toxin warhead, cleavage site, and half-life extender) of the original 15-mer. Minimization was performed in the same manner as described above, however for the sake of time, equilibration and production steps were not performed on the integrin structures.

## DECLARATIONS

No AI was used by the authors in the composition of this paper. All authors except JK are shareholders in Catessa Biotherapeutics. Data are available upon request.

## ACKNOWLEDGEMENTS

We thank Dr. Mark Norman for his input on Catamer design, and Drs. Songli Wang and Andrew Lezia for their valuable scientific feedback on the experimental design and the overall direction of this project.

## AUTHOR CONTRIBUTIONS

AK contributed to the first draft; AK, HX to the catamer concept and specific details of the molecular design; SX, RB and JC to the details of molecular design and chemistry; JK to execution of the molecular simulations. All authors contributed to revision of the manuscript.

## Notes

### Competing Interest Statement

The authors have declared no competing interest.

