## Supplementary Figures for "Catamers: Multi-specific therapeutics that concatenate individual warheads on a DNA scaffold via Watson-Crick interactions"

### SUPPLEMENTARY MATERIAL

**Fig. S2:** Toxin warhead (DXd) with PSMA cleavage site and other functionality used in the modeling.

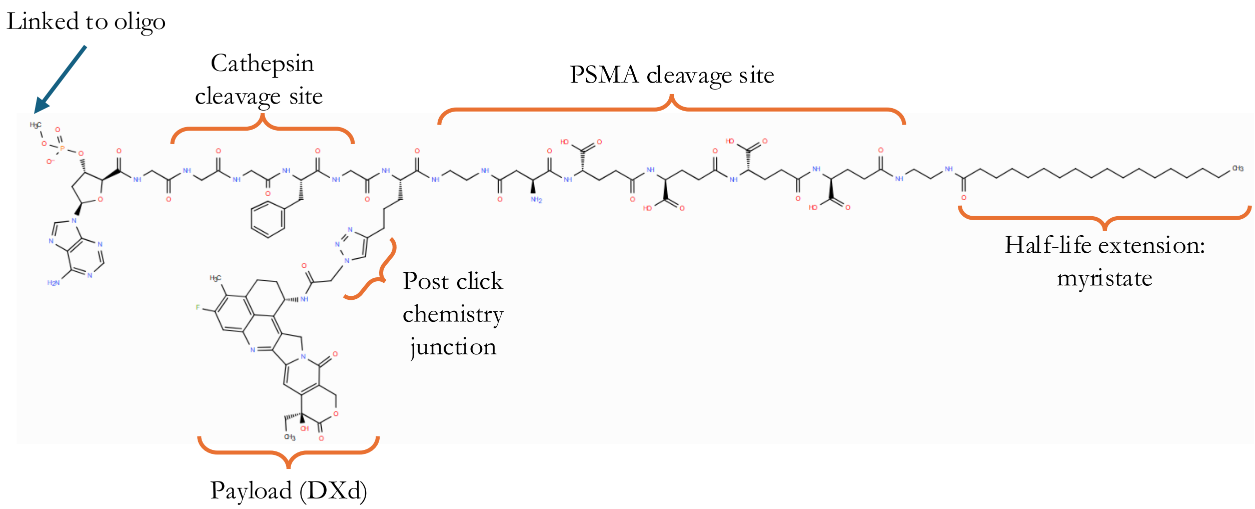

Fig. S3: Other models related to Fig. 3. A. Model with PSMA protease bound to cleavage site. The larger protease (brown) clashes with the bound HSA protein (purple). The grey portion of the complex is the PSMA protein that binds the PSMA-binding, tumor-targeting warhead. B. Model showing complex of HSA with FcRN (gold), with the catamer bound to HSA via myristate (see Fig. 3B for comparison without the FcRN protein bound).

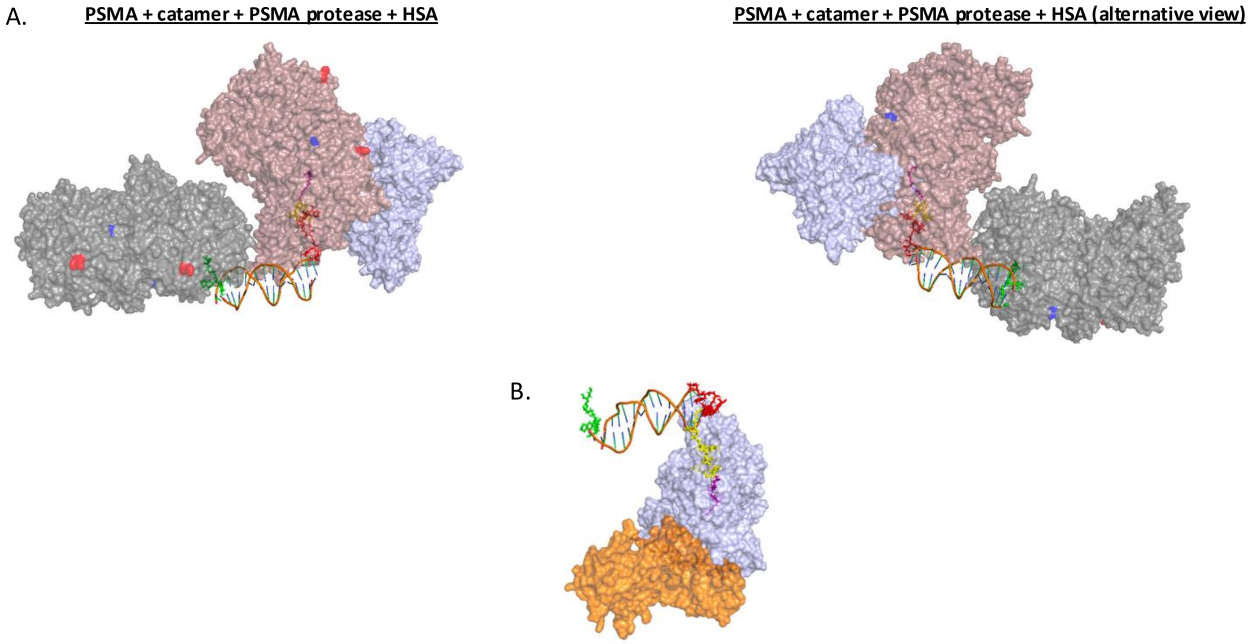

Fig. S4A Views of simulated catamer complexes. A. Alternative view of complex shown in Fig. 4B with 20 PEG units. B. Energy-minimized PSA-protease complex that contains catamer with a polyglycine linker (5 units) between the myristate and protease-cleavage sites. Coloring is as described in Fig. 3. C. PSMA protease complex (20 glycine units). The protease and HSA are separated by 5-6 Å in both B. and C.

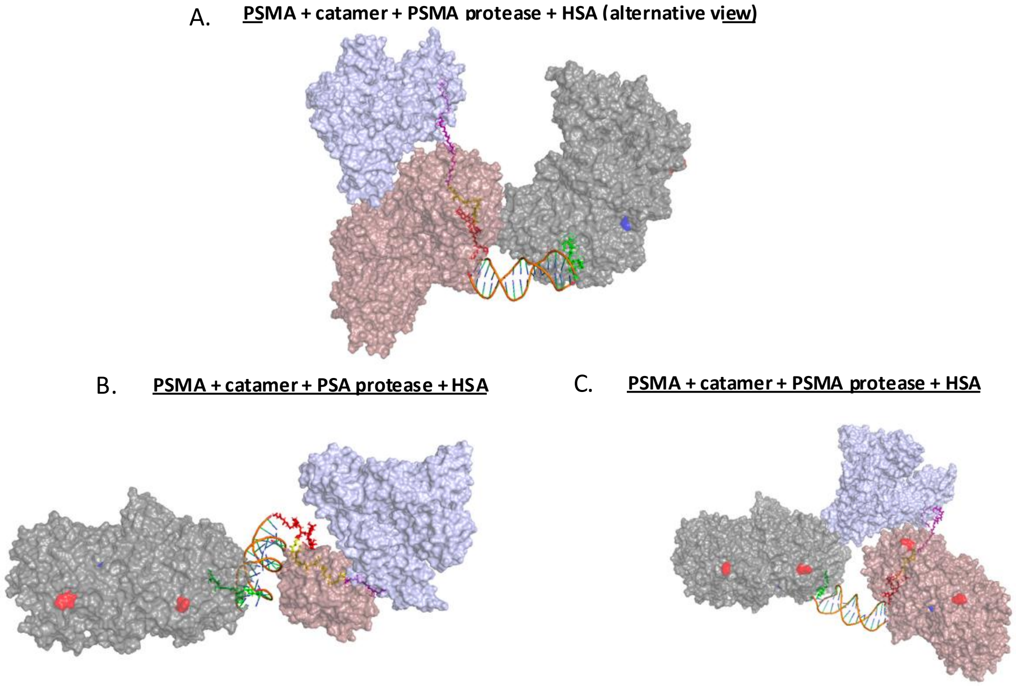

Table S4: General spacer types. The spacer is any chemical moiety interposed between a half-life extender (in the case described here, a myristate fatty acid moiety) and a peptide or peptide analog, which covalently links the fatty acid to the peptide and modulates spatial separation, flexibility, solubility, charge, or steric properties. The spacer may comprise amino acids, peptides, alkylene or heteroalkylene groups, polyether units, or combinations thereof, and may be cleavable or non-cleavable. The spacer may be attached through the N-terminus, C-terminus, or side-chain of the peptide. Note that spacers may be used in linkers to separate other moieties in the catamer. *OEG: Oligo(ethylene glycol), 2-10 units

| Spacer type | Examples |
|---|---|
| **Amino-Acid–Based Spacer** | Naturally occurring or non-naturally occurring amino acids, such as γ-Glu, Asp, Lys and side-chain conjugation |
| **Oligoether / PEG-Type Spacer** | PEG from 1 to about 20 repeating units |
| **Short Peptide / Oligopeptide Spacers** | Short peptide or peptide-like fragment, including dipeptides, tripeptides |
| **Aliphatic and Semi-Rigid Spacers** | Alkyl, aryl or heteroalkyl chain, may substituted and optionally containing amide, ester, urea, carbamate, or ether functionalities |
| **Hybrid Spacer** | γ-Glu–OEG*, Lys–PEG, multi-block linkers |

Fig. S5: Modeled superposition of integrin-binding peptide (A20FMDV2 ) and minibinder bound to αvβ6 integrin. The modeled peptide was fit into the minibinder pocket manually prior to energy minimization. White stick model is the minibinder; yellow space-filling structure is the peptide and the beige protein is the integrin. A. With C-terminus visible (red). B. With N-terminus visible (blue). Note integrins are type I membrane proteins. C. Another view of the structure shown in Fig. 5C that shows the catamer Format A structure clearly (see Fig. 5). The grey parallelogram represents the plane of the cell membrane predicted from the orientation of the PSMA and integrin termini.

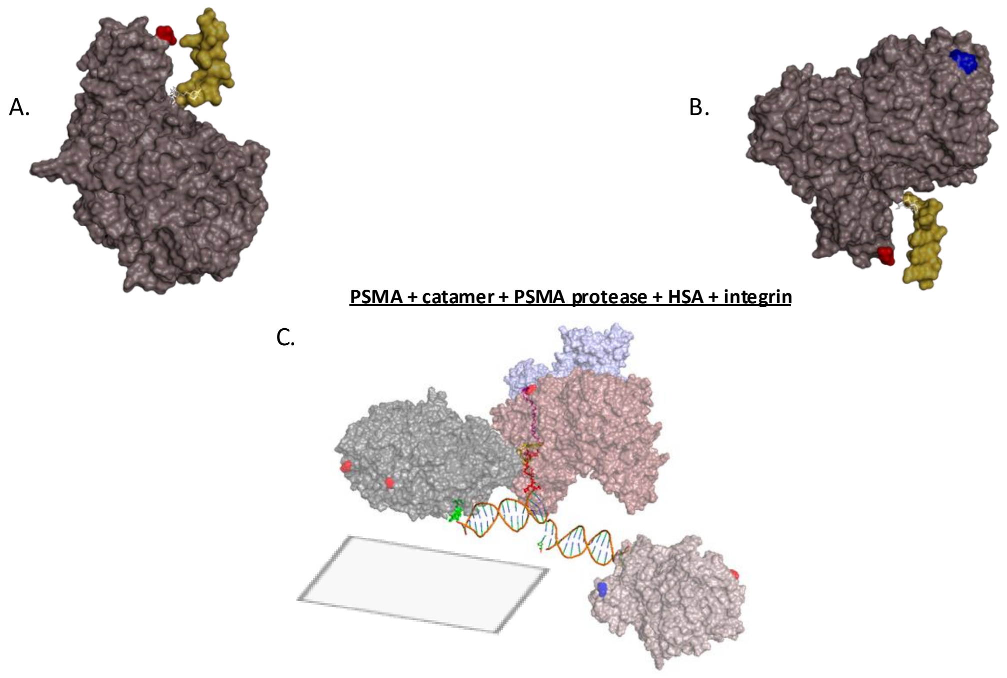
